# Isolated Mouse Adult Cardiomyocytes Display Minimal Mitochondrial ATP Demand and Maximal Reliance on Glycolysis

**DOI:** 10.1101/2024.11.17.623751

**Authors:** Ushodaya Mattam, Noble Kumar Talari, Ashish Rao Sathyanarayana, Sobuj Mia, James Frazier, Sam Slone, Konstantinos Drosatos, Karthickeyan Chella Krishnan

**Affiliations:** Department of Pharmacology, Physiology, and Neurobiology, University of Cincinnati College of Medicine, OH, USA; Department of Surgery, School of Medicine, Duke University, NC, USA

## Abstract

Mitochondrial maladaptation is a hallmark of heart failure, contributing to impaired energy production and contractile dysfunction. Understanding the bioenergetics of cardiomyocytes under healthy and pathological conditions is critical for characterizing mitochondrial maladaptation. While adult cardiomyocytes from rodents are a widely used model, recent studies have reported oligomycin insensitivity in these cells, a phenomenon often overlooked. This has led to incomplete assessments of key bioenergetic parameters, such as ATP-linked respiration and glycolytic capacity, focusing primarily on basal and maximal respiration. In this study, we performed a comprehensive characterization of bioenergetic and glycolytic profiles in three cardiomyocyte models: neonatal rat ventricular myocytes (NRVMs), human induced pluripotent stem cell-derived cardiomyocytes (hiPSC-CMs), and mouse adult cardiomyocytes (mouse ACMs). Our findings demonstrate distinct metabolic adaptations in mouse ACMs, revealing critical insights into mitochondrial function, ATP demand, and glycolytic reliance. These results underscore the importance of selecting appropriate cellular models for studying mitochondrial bioenergetics in cardiac physiology and pathology.

---

Mitochondrial maladaptation has been recognized as a pathological mechanism in the development of heart failure ^1,2^. The primary putative mechanism linking mitochondrial maladaptation to heart failure is reduced oxidative respiration leading to contractile failure. Therefore, mitochondria are an attractive target for heart failure therapy ^2^. To elucidate and characterize mitochondrial maladaptation in cardiomyocytes, one must compare cellular bioenergetics between healthy and heart failure conditions. For this purpose, investigators have optimized a standardized protocol ^3^ and have reported differential cellular bioenergetic profiles between sex ^4^ and ventricles ^5^. However, a notable and consistent finding across these studies, often overlooked or insufficiently explored, was that mouse adult cardiomyocytes (mouse ACMs) were unresponsive to oligomycin (ATP synthase inhibitor). As a result, these studies typically report only basal or maximal respiration rates, neglecting ATP-linked respiration and glycolytic capacity, thereby leaving important aspects of cellular bioenergetics unexplored.

To this end, we have comprehensively characterized the bioenergetic and glycolytic capacities of neonatal rat ventricular myocytes (NRVMs), human induced pluripotent stem cell derived cardiomyocytes (hiPSC-CMs) and ACMs and reported their oxygen consumption (OCR) and proton efflux rates (PER), respectively. We found that, unlike NRVMs and hiPSC-CMs, blocking ATP synthase by oligomycin in ACMs minimally decreased their basal OCR (**Figs. 1A – C**). Concomitantly, we found that oligomycin addition significantly increased the basal PER in both NRVMs and hiPSC-CMs as a compensatory mechanism for ATP production (**Figs. 1D – E**). In strong contrast, the basal PER was minimally increased in ACMs (**Fig. 1F**). Additionally, a notable observation in ACMs was the remarkable similarity between their OCR and PER profiles compared to both NRVMs and hiPSC-CMs (**Figs. 1C and F**).

**Figure 1.**
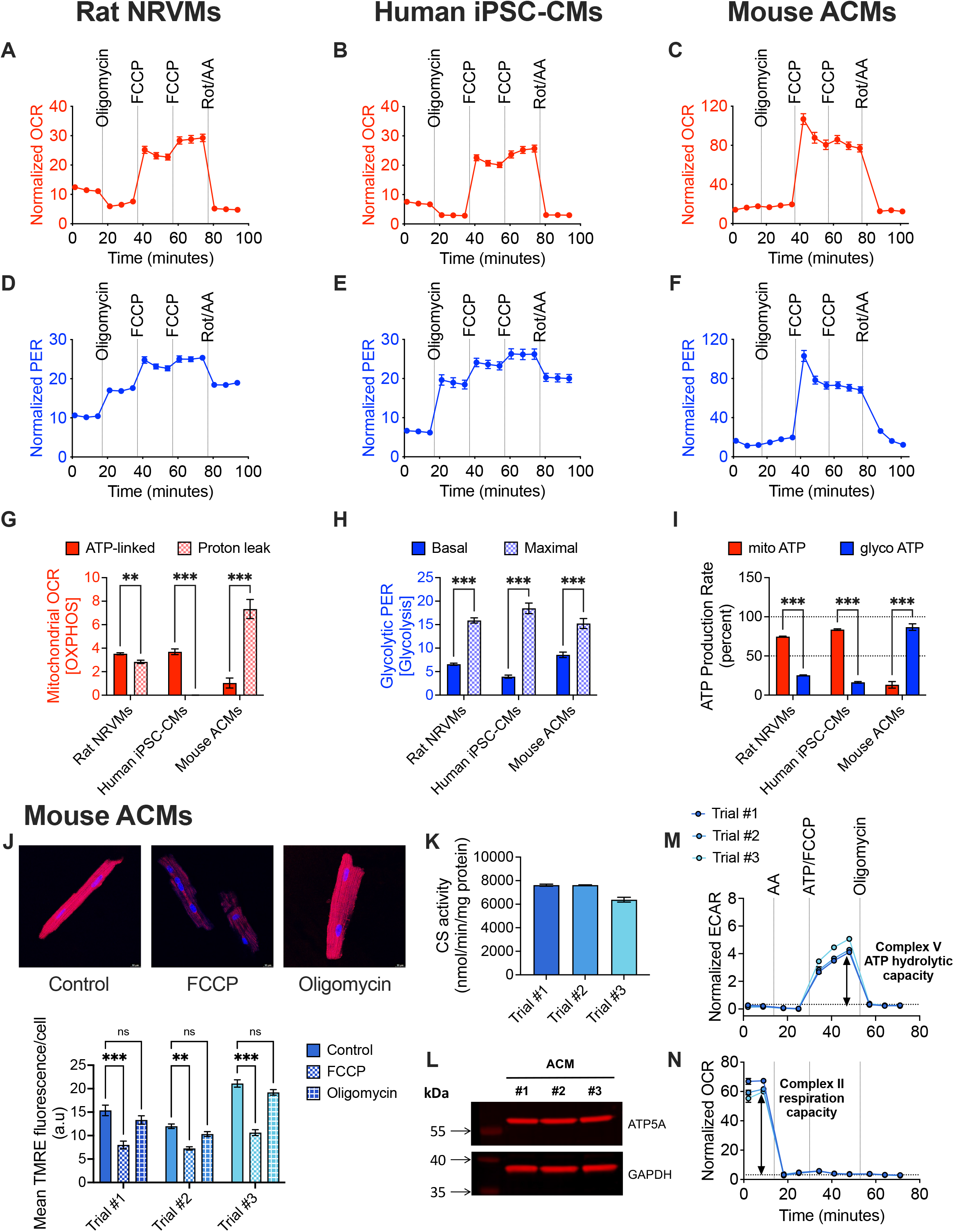
Isolated ACMs rely more on glycolysis than mitochondria for ATP production. Oxygen consumption (OCR) and proton efflux rate (PER) traces of intact (**A and D**) NRVMs (20,000 cells/well on gelatin coated XF 96-well plates), (**B and E**) hiPSC-CMs (20,000 cells/well on gelatin coated XF 96-well plates) and (**C and F**) ACMs isolated from 10-week old male C57BL/6J mouse (4,000 – 6000 cells/well on laminin coated XF 96-well plates) in basal and after sequential injections of oligomycin (NRVMs: 2 µM, iPSC-CMs and ACMs: 2.5 µM), FCCP (NRVMs, iPSC-CMs and ACMs: 1 µM), and rotenone/antimycin A (NRVMs: 0.5 µM, iPSC-CMs and ACMs: 1 µM). Measures were normalized by cell number counted using BioTek Cytation 5 Cell Imaging Multimode Reader. All intact bioenergetic assays were performed in XF DMEM or RPMI medium (pH 7.4) containing 10mM glucose, 1mM pyruvate, and 2mM glutamine. (**G**) ATP-linked (OCR: datapoint 6 subtracted from 3) and proton leak (OCR: datapoint 15 subtracted from 6) respiration; (**H**) basal (Glycolytic PER: datapoint 3) and maximal (Glycolytic PER: datapoint 6 subtracted from 3) glycolytic capacities; (**I**) mitochondrial and glycolytic ATP production rates from NRVMs (n = 5 replicates), hiPSC-CMs (n = 14 replicates) and ACMs (n = 8-16 replicates isolated from 4 mice on separate days). (J) (Top) Representative confocal images of ACMs stained with TMRE under control conditions and following treatment with 1µM FCCP or 2.5µM oligomycin. Scale bar = 10 µm. (Bottom) TMRE fluorescence was captured using excitation at 548 nm and emission at 574 nm. Mean TMRE fluorescence per cell was quantified using the ImageJ software (n = 6-24 individual cells isolated from 3 mice on separate days). (K) Citrate synthase (CS) activity measured using 1.25 µg of frozen whole-cell lysates prepared from ACMs (n = frozen ACMs from 3 mice). Reaction mixtures included the sample, 100 µM DTNB and 0.3 mM Acetyl-CoA, to which 0.5 mM oxaloacetic acid was added and the increase in absorbance at 412 nm was measured. Enzyme activities were normalized to protein levels. (L) Immunoblot analyses of mitochondrial complex V subunit protein, ATP5A, in ACMs (n = frozen ACMs from 3 mice). GAPDH was used as a loading control. (M) Extracellular acidification (ECAR) and (**N**) oxygen consumption rate (OCR) traces from 5µg of frozen whole-cell lysates prepared from ACMs (n = frozen ACMs from 3 mice) sustained by 5mM succinate/2µM rotenone before and after sequential injections of 2µM antimycin A, 20 mM ATP/1µM FCCP, and 5µM oligomycin. Measures were normalized to protein levels. The complex V ATP hydrolytic capacity was quantified as the difference in acidification (ECAR) rates between ATP/FCCP and oligomycin injections, while the complex II respiration capacity was quantified as the difference in respiration (OCR) rates before and after antimycin A injection. All frozen bioenergetic assays were performed in MAS buffer (70mM sucrose, 220mM mannitol, 10mM KH_2_PO_4_, 5mM MgCl_2_, 1mM EGTA, and 2mM HEPES, pH 7.2) containing 5mM succinate, 2µM rotenone and 100 µg/ml of cytochrome c. Data are presented as mean ± SEM. P values were calculated by (**G – I**) multiple t tests or (**J**) one-factor ANOVA, corrected by post hoc ‘Holm-Sidak’s’ multiple comparisons test. ns, non-significant; **P < 0.01; ***P <0 .001.

Follow-up analyses revealed that ATP-linked (oligomycin sensitive) and proton leak (oligomycin insensitive) respiration rates were remarkably different among the three cardiomyocyte populations. Specifically, we observed that proton leak was significantly higher than ATP-linked respiration in ACMs, while ATP-linked respiration was higher in both NRVMs and hiPSC-CMs (**Fig. 1G**). FCCP is a mitochondrial uncoupler that mimics energy demand by operating the electron transport chain at maximal capacity *via* mitochondrial membrane potential depletion. FCCP stimulation of respiration in all three cardiomyocytes highlights that the oligomycin insensitivity observed in ACMs was due to reduced mitochondrial ATP demand and not cell viability (**Figs. 1A – C**). Since ACMs did not depend on mitochondria for ATP, we reasoned that they may depend on alternative pathways such as glycolysis for ATP production. To test this, we calculated the basal and oligomycin-induced maximal glycolytic capacities and found that basal glycolysis was higher and more than 50 percent of their maximal capacity in ACMs (**Fig. 1H**), suggesting ACMs are already relying maximally on glycolysis. Additionally, we found that glycolytic ATP production rate was significantly greater than mitochondrial-dependent ATP production rate in ACMs, which was the complete opposite of what was observed in both NRVMs and hiPSC-CMs (**Fig. 1I**). Taken together, our findings indicate that isolated ACMs rely more on glycolysis than mitochondria for ATP production.

Several factors influence oligomycin sensitivity, including proton leak, ATP demand, ATP synthesis rate, defects in substrate supply or electron transport chain activities. Here, we have systematically examined this and provided evidence that reduced mitochondrial ATP demand in ACMs might be the fundamental reason for oligomycin insensitivity. Since proton leak is often associated with reduction in mitochondrial membrane potential (ΔΨm), we monitored ΔΨm in ACMs by TMRE fluorescence. As expected, compared to the control, oligomycin addition did not change ΔΨm suggesting oligomycin insensitivity. However, FCCP-induced membrane depolarization significantly reduced ΔΨm (**Fig. 1J**). Next, we examined the activity of TCA cycle regulatory enzyme, citrate synthase (CS), as a surrogate for TCA cycle flux, and found that ACMs show robust CS activity (**Fig. 1K**). Our immunoblot analyses also revealed no defects in mitochondrial complex V subunit levels in ACMs (**Fig. 1L**). We next measured the mitochondrial ATP hydrolytic capacity (reverse ATP synthase activity) of ACMs and observed substantial activity, which was completely inhibited following oligomycin injection (**Fig. 1M**). Finally, we showed that ACMs have significant succinate-driven complex II respiration capacity that was inhibited by antimycin A (complex III inhibitor) emphasizing there were no defects in substrate supply or electron transport chain activities (**Fig. 1N**).

In summary, our findings demonstrate that oligomycin insensitivity in ACMs cannot be attributed to defects in substrate supply, electron transport chain, or ATP synthase activity. Instead, our data clearly establish that ACMs possess a fully functional and oligomycin-sensitive mitochondrial ATP synthase, which remains underutilized for coupled respiration due to reduced mitochondrial ATP demand. One possible explanation for this phenomenon could be the extreme ‘ischemic’ stress experienced by the ACMs during isolation and/or their adaptation from the *in vivo* to *in vitro* system. Based on these findings, we propose that ACMs are more suitable for assessing individual electron transport chain activities, whereas hiPSC-CMs are better suited for evaluating coupled respiration. Ultimately, our study underscores the importance of careful interpretation of cellular bioenergetics in heart failure research.

## ACKNOWLEDGEMENTS

We thank all the members of the K lab for their assistance and helpful discussion. We also thank the Live Microscopy Core for their assistance, especially Chet T. Closson. We would like to acknowledge Drs. Michael Tranter and Hong-Sheng Wang for their in-kind donations of NRVMs and hiPSC-CMs, respectively.

## AUTHOR CONTRIBUTIONS

K.C.K. conceived the study. K.C.K., A.R.S., N.K.T., and U.M. designed, performed experiments, or analyzed the data. J.F. assisted in seahorse experiments. S.S. performed NRVM isolations. S.M., and K.D. performed or supervised mouse adult cardiomyocyte isolations. K.C.K., U.M., and N.KT. drafted the manuscript, and all authors read or revised the manuscript.

## FUNDING INFORMATION

This work was supported by R01 HL167670 (K.C.K) and P30 DK078392 (Live Microscopy Core) of the Digestive Diseases Research Core Center in Cincinnati. The funders had no role in study design, data collection and interpretation, or the decision to submit the work for publication.

## CONFLICTS OF INTEREST

All authors declare no conflict of interests.

## Notes

### Competing Interest Statement

The authors have declared no competing interest.

